# CTE-Type Tau Filaments in Alzheimer’s Disease with Co-morbid LATE-NC

**DOI:** 10.1101/2025.10.21.683735

**Authors:** Jaimin K. Rana, Emile S. Pinarbasi, Martin G. Fernandez, Vikas Navratna, Kyle S. Conway, Andrew P. Lieberman, Sami J. Barmada, Shyamal Mosalaganti

## Abstract

Alzheimer’s disease (AD), the most common neurodegenerative disease, is defined by β-amyloid plaques and tau neurofibrillary tangles. Tau filaments in AD adopt the “Alzheimer’s fold”, which is distinct from other tauopathies and highly conserved across sporadic and familial AD. However, to date, structural studies have focused on pure AD, despite the high prevalence of comorbid pathologies. In particular, up to half of AD patients harbor limbic-predominant age-related TDP-43 encephalopathy neuropathologic change (LATE-NC). This co-pathology— consisting of mislocalized and aggregated TDP-43, mainly restricted to the medial temporal lobe—is associated with accelerated clinical decline and more severe tau pathology. Whether there is a structural basis for this clinical synergy remains unknown. Here, using cryo-electron microscopy, we determine the structure of tau filaments from three AD patients with LATE-NC. We show that in addition to the expected Alzheimer’s fold tau filaments, all cases exhibit a distinct fibril morphology identical to chronic traumatic encephalopathy (CTE) fold tau. Additional sampling revealed CTE neuropathology in one patient, suggestive features without definitive CTE in the second, and no evidence of CTE in the third. These findings raise questions about the relationship between LATE-NC and CTE, the effect of TDP-43 on tau conformation, and the etiology of LATE-NC.

## Introduction

Alzheimer’s disease (AD) is the most common neurodegenerative disease, affecting approximately 7.2 million people in the United States alone^1^. Neuropathologically, AD is characterized by extracellular amyloid β (Aβ) plaques and intracellular neurofibrillary tangles composed of protein tau^2^.

AD, together with familial British dementia (FBD), familial Danish dementia (FDD), primary age-related tauopathy (PART), and chronic traumatic encephalopathy (CTE), belongs to a subset of mixed 3R+4R tauopathies in which tau aggregates are composed of isoforms containing three (3R: R1, R3, R4) as well as four (4R: R1–R4) repeats of the microtubule-binding domain^3–6^. This subset of tauopathies can be further classified based on distinct ultrastructural features of tau^3^. Cryo-electron microscopy (cryo-EM) analysis of patient-derived pathological aggregates revealed a robust cross-β architecture for tau fibrils, formed by the in-register stacking of β-strands within the microtubule-binding repeat domains^7^. Tau filaments in AD, PART, FBD, and FDD contain two different structural polymorphs: paired helical filament (PHF) or straight filament (SF), where two tau protofilaments form an interface mediated by amino acids 332-336 (PHF) or 317-324 and 312-321 (SF). The structural signatures of PHF and SF polymorphs— collectively referred to as the Alzheimer’s fold or AD-fold—are highly conserved across sporadic and familial AD cases^3,7–9^. By contrast, the architecture of tau filaments in patients with CTE is distinct. CTE-fold tau protofilament still adopts the same overall ‘C-shaped’ morphology, but the intra-filament interface is formed by amino acids 324-329 (type I) or 331-338 (type II)^10^, resulting in a more open conformation in comparison to PHF or SF. In addition to CTE, this unique tau conformation has also been observed in subacute sclerosing panencephalitis (SSPE) and ALS/PDC (amyotrophic lateral sclerosis / Parkinsonism-dementia complex of Guam)^10–12^, neurodegenerative conditions that, like CTE, are marked by co-pathology involving the RNA-binding protein TDP-43^13,14^.

To date, structural characterization of pathological aggregates in AD has been restricted to “pure AD” cases without taking into account co-morbid pathologies that are increasingly recognized as highly prevalent, especially in the elderly^15–17^. In particular, up to 50% of patients with AD also display limbic predominant age-related TDP-43 encephalopathy neuropathologic change (LATE-NC)^18–20^. LATE-NC is characterized by the accumulation of mislocalized and aggregated TDP-43, predominantly in medial temporal structures including the amygdala and hippocampus^21^, and is independently associated with amnestic dementia^22–24^. Furthermore, AD patients with co-morbid LATE-NC exhibit more widespread distribution of tau pathology, higher Braak stage, and more severe clinical decline^18,20,25,26^, indicating synergy between AD and LATE-NC with significant pathological as well as clinical consequences. Even so, whether the presence of co-aggregating proteins such as TDP-43 influences tau architecture in these diseases remains unknown.

Here, we show that co-morbid AD with LATE-NC (AD+LATE-NC) is a mixed pathology characterized by pathogenic accumulations of tau, Aβ, TDP-43, and TMEM106B. We provide the first comprehensive structural landscape of tau filaments from the amygdala of three AD+LATE-NC patients, uncovering CTE-fold tau in addition to classical AD-fold tau in all three cases. Moreover, through extensive neuropathological sampling, we detected pathognomonic CTE lesions and suggestive features of CTE in some instances, despite the apparent absence of head trauma or injury. Collectively, our results provide original structural evidence for the coexistence of different tau ultrastructural polymorphs in disorders of mixed pathologies. In addition, our data support emerging evidence for the accumulation of CTE-fold tau in diseases associated with TDP-43 neuropathology.

## Results

### Multiple insoluble proteins are present in the amygdala of a patient with AD+LATE-NC

We selected a patient with AD and co-morbid LATE-NC for structural analysis (see **Tables 1 and 4** *for further clinical and neuropathologic details)*. Tau immunostaining in the amygdala highlighted dense tau pathology composed predominantly of NFTs and neuropil threads. Additionally, frequent diffuse and neuritic amyloid plaques were observed by amyloid-β (Aβ) immunostaining (**Fig. 1a**, right panel). Dense phosphoTDP-43 (pTDP-43) pathology, consisting of “NFT-like” neuronal cytoplasmic inclusions in the entorhinal cortex (“Pattern β^27^)”, confirmed the presence of LATE-NC (**Fig. 1a**, middle panel). To explore the molecular content of these inclusions, we extracted sarkosyl-insoluble material from the amygdala, a region showing prominent neuropathology in both AD and LATE-NC. The resulting samples were highly enriched for high molecular weight species of phosphorylated tau (pTau), Aβ, and pTDP-43 (**Extended Data Fig. 1)**. Consistent with the accumulation of TMEM106B in age-related diseases, we also observed TMEM106B in negative stains of detergent-insoluble material from AD+LATE-NC (**Fig. 1b**, right panel). Finally, we confirmed distinct filaments comprised of pTau, pTDP-43, and TMEM106B by immuno-electron microscopy (immuno-EM) using antibodies specific for each protein (**Fig. 1b**; **Table 2**).

**Figure 1:**
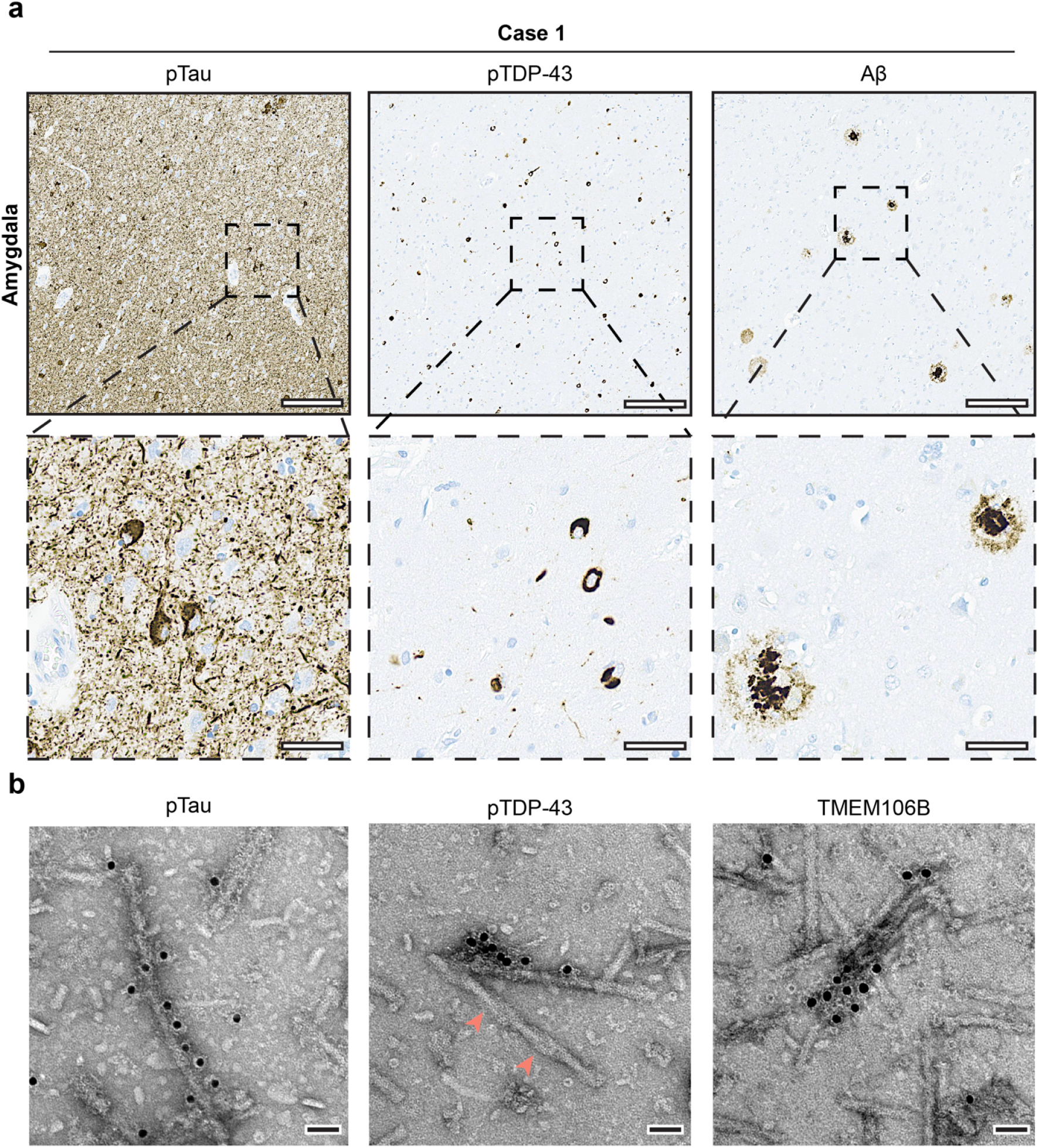
Detergent insoluble material from Alzheimer’s disease (AD) with co-morbid limbic age-related TDP-43 encephalopathy neuropathological change (LATE-NC) displays several distinct fibril types. **a**. Representative images of amygdala from case 1 immunostained with AT8 (specific for pS202 and T205 tau; pTau, top left), anti-pS409/S410 TDP-43 antibody (pTDP-43, middle), and anti-amyloid beta antibody (Aβ, top right). Insets show high magnification views. Scale bars: 300 µm (top) and 60 µm (bottom). **b**. Representative negative stain EM images of immunogold-labeled sarkosyl-insoluble material (anti-pTau, left; anti-pTDP-43, middle; anti-TMEM106B, right) from Case 1. Arrowheads (warm pink, middle) point to a tau PHF fibril as evident from the characteristic shape^7^ that is not recognized by the anti-pTDP43 antibody, highlighting the selectivity of immunogold labeling. Scale bar: 50 nm.

### Tau filaments in AD+LATE-NC display both AD- and CTE-fold conformations

Cryo-electron microscopy (EM) of sarkosyl-insoluble material from AD+LATE-NC amygdala revealed an ensemble of filaments demonstrating varying thickness and crossovers distances, suggesting the presence of multiple morphologies (**Extended Data Fig. 2a**). We collected 18,070 images and extracted 2,374,377 fibril segments. 2D averaging, followed by selecting particles based on dendrogram separation and subsequent 3D reconstruction, enabled us to determine the structures of SF and PHF tau at 2.7 and 2.9 Å, respectively (**Fig. 2a, b, Extended Data Fig. 2b and 3a; Table 3**). These structures showed no deviation from previously reported tau structures in AD patients, providing additional support for the conserved signature of AD-fold tau even in the presence of mixed pathologies. SF and PHF tau in AD+LATE-NC are composed of two ‘C-shaped’ protofilaments with distinct interfaces^7,9^. The resolution of our map enabled us to build *de novo* models. Additionally, we fit the published Protein Data Bank (PDB) models of SF (5O3L) and PHF (5O3T)^7^ tau and used real-space refinement in PHENIX^28^. Both approaches yielded models that were nearly identical to previously published models of SF and PHF, with a mean RMSD of < 1 Å (**Extended Data Fig. 2c, 2d**). Each protofilament showed precise overlap with V306 to F378 core residues of 4R tau from previously reported structures, with sequences beyond this region being too flexible to resolve. Our cryo-EM maps also showed additional densities surrounding the SF and PHF protofilaments, as previously observed in the structures of SF and PHF in pure AD cases (**Fig. 2a, b**)^7^. Given the similarity between *de novo* models and previously deposited ones, we chose not to deposit additional entries into the PDB.

**Figure 2:**
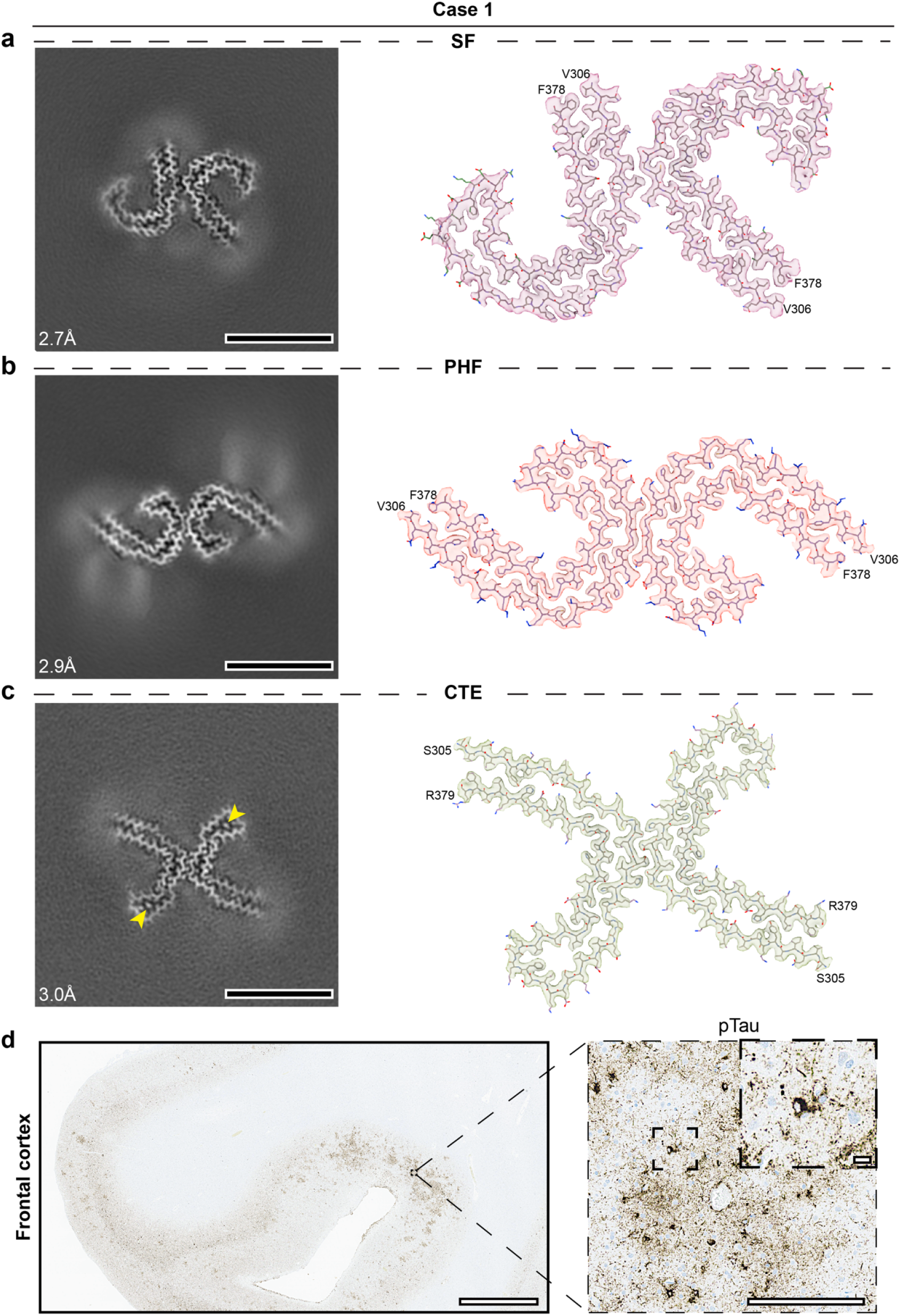
High-resolution structural analysis uncovers three separate tau morphologies in AD+LATE-NC. **a-c**. Structures of sarkosyl-insoluble tau filaments isolated from the amygdala of case 1. Cross-sections through the cryo-EM reconstructions, perpendicular to the helical axis (left), and the cryo-EM density map with a fit of the tau atomic model (right) of (a) straight filaments (SF), (b) paired helical filaments (PHF), and (c) chronic traumatic encephalopathy (CTE) fold tau. The cross-sections are approximately one rung thick. Resolutions of structures were calculated from the gold-standard Fourier shell correlation between two half-maps. Scale bar: 10 nm. Yellow arrowheads highlight non-proteinaceous density at the center of the β-helix turn in CTE-fold tau. Cryo-EM cross-sectional densities for two identical protofilaments fitted to tau amino acids F378 to V306 for SF (magenta, PDB: 5O3T) and PHF (warm pink, PDB: 5O3L), and R379 to S305 for CTE (light green, PDB: 6NWP). **d**. CTE lesion in the frontal cortex immunostained for pTau. Insets show high magnification views. Scale bars: 3 mm (left) and 200 µm (insets).

In addition to AD-fold tau, we noticed distinct class averages that resembled CTE-fold tau^10^ in the initial reference-free 2D classification (**Extended Data Fig. 2b**). We could resolve the structure of these filaments at a nominal resolution of 3.0 Å (**Fig. 2c, Extended Data Fig. 2b and 3a; Table 3)**, confirming the presence of type I CTE-fold tau. These filaments consist of two protofilaments adopting a more open β-helix turn conformation in comparison to PHF or SF tau^10^. We were able to unambiguously fit the previously published PDB model for CTE type I (6WNP), consisting of two distinct protofilaments, each made up of core residues extending from S304-R379 in our structure. We also observed a non-proteinaceous density at the center of the β-helix turn in our CTE-fold tau map, analogous to the previously reported structure (**Fig. 2c**, yellow arrowheads). Although previous investigations of clinically diagnosed CTE cases demonstrated two types of tau filament morphologies (type I and II), we were unable to identify filaments corresponding to CTE type II in our case of AD+LATE-NC. This is consistent with the predominance of type I over type II fibers in CTE, potentially owing to the relatively weak interaction at the type II protomer interface^10^.

Together, these results suggest that tau filaments exist as SF, PHF, and CTE type I structural polymorphs in AD+LATE-NC.

### Neuropathologic characterization reveals CTE in a patient with AD+LATE-NC

CTE is a neurodegenerative disease tightly associated with repetitive head injury, particularly in the context of contact sports^29–31^. It is neuropathologically defined by “pathognomonic” lesions of neuronal tau aggregates arranged around small vessels in the depths of cortical sulci^32^. Based on our structural findings indicating the presence of CTE-fold tau, we performed additional cortical sampling per consensus guidelines to evaluate for undiagnosed CTE pathology in our case of AD+LATE-NC^32^. In doing so, we observed a single pathognomonic CTE lesion in the frontal cortex (**Fig. 2D**), consistent with previously unappreciated CTE in this individual.

TDP-43 pathology occurs in ∼40% of CTE cases^33^, but unlike LATE-NC, it can extend to involve the spinal cord and other regions outside of the medial temporal cortex^34,35^. Broad cortical sampling in this patient uncovered rare pTDP-43 deposits and neurites without NCIs in the frontal cortex, and no pTDP-43 reactivity in the brainstem or spinal cord, supporting the neuropathological diagnosis of LATE-NC (**Table 4**). Furthermore, a comprehensive review of the patient’s clinical records and obituary did not reveal any history of traumatic head injury or contact sport participation^29–31^.

### CTE-fold tau is a consistent feature of AD+LATE-NC

To determine whether the presence of CTE-fold tau and neuropathologic CTE in AD+LATE-NC reflected a fundamental relationship between the two disorders, we selected two additional AD+LATE-NC cases for detailed structural analyses.

Case 2 exhibited high-stage LATE-NC in addition to high AD (**Tables 1 and 5**). Aging-related tau astrogliopathy (ARTAG^36^) was also noted in the medial temporal lobe, lobar, subcortical, and brainstem regions. While the majority of ARTAG was limited to white matter in these regions, we also observed subpial, subependymal, perivascular, and gray matter ARTAG (see **Table 5** for additional details). Tau immunostaining demonstrated dense tau pathology in the amygdala, with NFTs, neuropil threads, and astrocytic tau deposits (**Extended Data Fig. 4**). Moderate diffuse and neuritic plaques in the same region were detected via Aβ immunostaining (**Extended Data Fig. 4**). Staining for pTDP-43 highlighted dense pathology, predominantly round NCIs and short neurites in the basolateral nuclear group, consistent with LATE-NC pattern L^27^ (**Extended Data Fig. 4**).

Case 3 likewise showed high-stage LATE-NC but only intermediate AD (**Tables 1 and 6**). The amygdala displayed moderate tau pathology, composed of NFTs, neuropil threads, and astrocytic tau (**Extended Data Fig. 5**). Aβ was absent from this region (**Extended Data Fig. 5**), but there was moderate pTDP-43 pathology composed of round NCIs and short neurites in the basolateral nuclear group, indicating LATE-NC pattern L^27^ (**Extended Data Fig. 5**).

Negative staining of the sarkosyl-insoluble fractions from cases 2 and 3 demonstrated a mixed population of filaments, similar to those found in case 1. Based on the characteristic width and crossover, we identified these as PHF and SF tau, as well as singlet and doublet species of TMEM106B (data not shown). We used cryo-EM to elucidate the structure of sarkosyl-insoluble filaments from the amygdala of cases 2 and 3. As with case 1, we detected three separate filament morphologies for tau, including SF, PHF, and CTE-fold type I (**Fig. 3**). We determined high-resolution structures for each species, ranging from 3.0-3.1 Å in case 2, and 3.2-3.4 Å in case 3 (**Fig. 3 and Extended Data Fig. 3b, c**). In keeping with our results from case 1, as well as previous studies^10^, we detected a prominent density at the core of the β-helix turn for CTE-fold type I, as well as additional densities bordering all three filament types (**Fig. 3**). Even after exhaustive classification on the entire dataset, we were unable to identify filament class averages resembling CTE-fold type II filaments. During our cryo-EM analysis, we observed that the proportion of SF tau outnumbered PHF tau in 2/3 cases, and the proportion of SF was significantly higher in the third case, in contrast to the previously reported ratio favoring PHF tau in other tauopathies, including pure AD (**Fig. 4a**)^7,9,37^.

**Figure 3:**
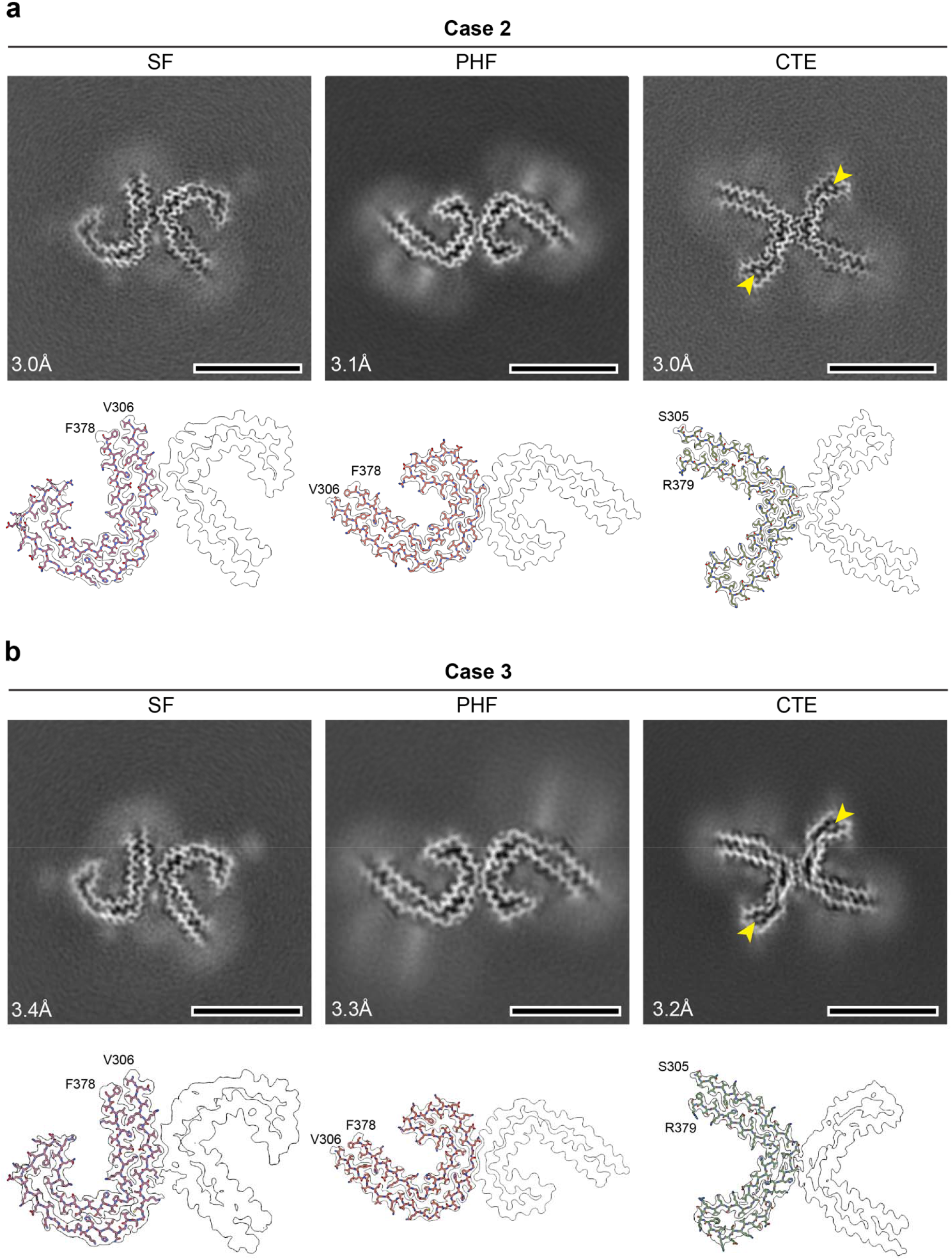
Structural evidence supports consistent deposition of CTE-fold tau in AD+LATE-NC amygdala. **a, b**. Structures of sarkosyl-insoluble tau filaments isolated from the amygdala of case 2 (a) and case 3 (b). Cross-sections through the cryo-EM reconstructions, perpendicular to the helical axis (top), and the cryo-EM density map with a fit of the tau atomic model (bottom) of SF (left), PHF (middle), and CTE (right). The cross-sections are approximately one rung thick. Resolutions of structures were calculated from the gold-standard Fourier shell correlation between two half-maps. Yellow arrowheads highlight non-proteinaceous density at the center of the β-helix turn in CTE-fold tau. Scale bar: 10 nm. Bottom panels: Cryo-EM cross-sectional densities for two identical protofilaments fitted to tau amino acids F378 to V306 for SF (PDB: 5O3T) and PHF (PDB: 5O3L), and R379 to S305 for CTE (PDB: 6NWP).

**Figure 4:**
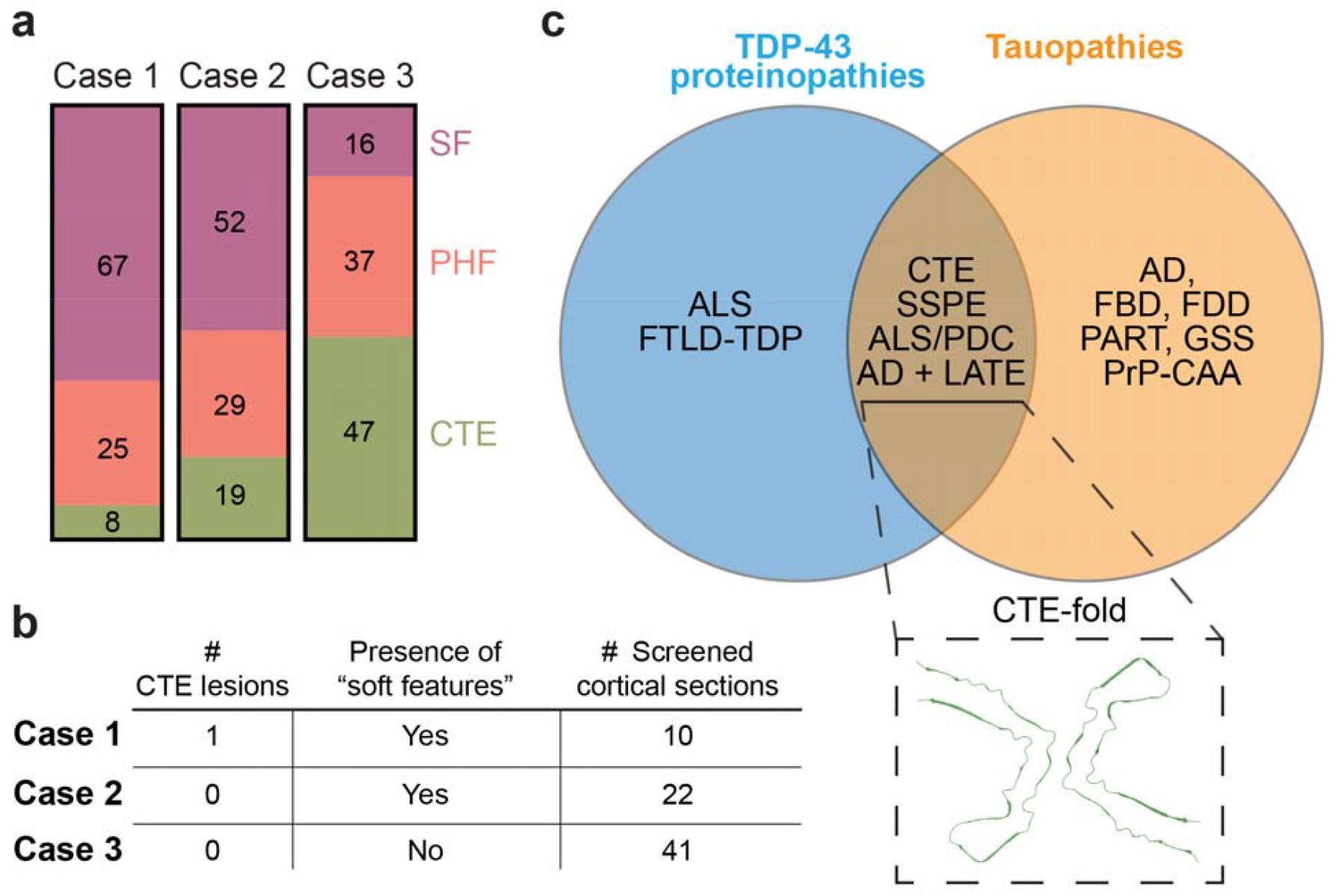
Tau adopts a CTE-fold in TDP-43 proteinopathies. **a**. Stacked bar graph depicting the percentage of tau structural polymorphs in AD+LATE Cases 1-3, as determined by cryo-EM analysis. **b**. Summary of neuropathologic features of CTE in the three cases. Surprisingly, the case with no detectable CTE pathology in the frontal cortex (case 3) shows the highest amount of CTE-fold tau. **c**. Schematic showing proteinopathies with TDP-43 or tau filament formation, specifically for conditions displaying the AD (SF and PHF) fold. In co-morbid mixed pathologies, marked by the presence of both TDP-43 and tau filaments, tau adopts the CTE fold (inset) to varying extent. Examples of such diseases that lie at this intersection include chronic traumatic encephalopathy (CTE), subacute sclerosing panencephalitis (SSPE), amyotrophic lateral sclerosis/parkinsonism-dementia complex of Guam (ALS/PDC), and limbic age-related TDP-43 encephalopathy with Alzheimer’s disease neuropathologic change (AD + LATE-NC). Abbreviations: FBD (familial British dementia), FDD (familial Danish dementia), PART (primary age-related tauopathy), GSS (Gerstmann-Sträussler-Scheinker disease), PrP-CAA (prion protein cerebral amyloid angiopathy), amyotrophic lateral sclerosis (ALS), frontotemporal lobar degeneration with TDP-43 pathology (FTLD-TDP).

### CTE neuropathology is inconsistently associated with AD+LATE-NC

Our discovery of CTE-fold tau in AD+LATE-NC in case 1 prompted a more extensive sampling of cortical tissue in the same case, revealing neuropathological evidence for CTE that had previously gone unappreciated. This observation, together with the presence of CTE-fold tau in two additional AD+LATE-NC cases, and the inherent challenges in recognizing CTE neuropathology among co-morbidities, motivated a comprehensive search for subtle as well as pathognomonic neuropathological signs of CTE in cases 2 and 3 [22 total sections in case 2, 41 in case 3 (**Fig. 4b**)]. In case 2, interpretation was limited due to the prominent, widespread ARTAG throughout the cortex (**Extended Data Fig. 6a-b, Table 4**). We observed a single lesion in case 2 that was highly suspicious for CTE (**Extended Data Fig. 6c**), with loosely clustered tau immunoreactivity at the depth of a sulcus. Closer inspection of these clusters revealed that they represent either the cores of neuritic plaques (**Extended Data Fig. 6c inset 1**) or astrocytic tau (**Extended Data Fig. S6c inset 2**). We also observed several features suggestive of CTE, including sulcal accentuation of astrocytic tau or tau neuropil staining (**Extended Data Fig. S6b, d**), without definite neuronal accumulation. Despite evaluation by a panel of neuropathologists, no consensus could be reached regarding these lesions and whether they met the threshold for diagnosing CTE. Therefore, we interpreted case 2 as possible CTE without a definitive lesion. In contrast, although half of the total tau filaments in case 3 were CTE-fold tau, a meticulous neuropathological review failed to detect signs of CTE in this patient (**Extended Data Fig. 7; Fig. 4b**). Once again, we extensively reviewed medical records, obituaries, and public records for these individuals but found no documented history of traumatic head injury or participation in contact sports.

## Discussion

Co-morbid neuropathologies are common, found in up to 80% of neurodegenerative disease post-mortem samples^38,39^. Even so, the large majority of studies focusing on protein conformations and their relationship to disease involve “pure” cases that exhibit only a single neuropathology. While such a focus effectively simplifies the isolation and analysis of disease-associated protein structures, it also ignores potential interactions between and among aggregate-prone proteins, and the cooperative effect these interactions may have on protein misfolding. Our analysis of AD cases with co-morbid LATE-NC, a neuropathological condition characterized by TDP-43 inclusions within the medial temporal lobe, uncovered CTE-fold tau in addition to the expected AD-fold tau common to AD and other tauopathies. Together, these observations add LATE-NC to a growing list of conditions involving both TDP-43 pathology and CTE-fold tau (**Fig. 4c**), suggesting critical interactions among these proteins that drive misfolding and aggregation in disease.

In all three cases of AD+LATE-NC that we sampled, the majority of tau filaments were classified as SF or PHF tau^7^ (**Figs. 2 and 3**). However, we observed an unprecedentedly higher proportion of SFs across all three cases. PHFs predominate (ratio of SF:PHF; 1:4) in published AD datasets, including those derived from limbic and subcortical brain regions^7,9^. SF assembly requires ubiquitination at lysine residues 317/321, representing the interface between filaments^37^, while PHFs rely on antiparallel stacking of residues 332-336^7,9^, suggesting that tau post-translational modifications (PTMs) may dictate the balance between SF and PHF tau. One possibility is that TDP-43 pathology—through loss of TDP-43 splicing activity or cytosolic sequestration of kinases^40–42^—modulates tau PTMs and thereby influences the SF:PHF ratio.

In addition to AD-fold tau, we found that all three individuals harbored CTE-fold tau, the structural polymorph first identified in CTE, which has never been previously reported as a component of AD^3^. Both AD- and CTE-tau folds exhibit two protofilaments with different interface residues. Time-resolved cryo-EM of *in vitro* PHF and CTE assembly reactions highlighted a first intermediate amyloid (FIA) common to both conformations that subsequently diverges and adopts distinct final structures after passing through several intermediate states^43^. A critical difference lies in the residue S324, which is buried within the protofilament interface in CTE-fold type I, whereas it is exposed and phosphorylated in AD-fold tau^37,44^. This event may preclude interactions among protofilaments required for assuming the CTE-fold tau conformation^10^.

The abundance of CTE-fold tau varied from 8% (Case 1) to almost 50% of the sampled tau filaments (Case 3**; Fig. 4a**). Strikingly, however, the proportion of CTE-fold tau showed little correlation with neuropathologic features of CTE. These results have at least two important implications: first, the formation of CTE-fold tau may not trigger CTE neuropathology in all cases, or the pathological changes indicative of CTE may occur in a delayed fashion following tau misfolding. Second, as discussed below, the limited tools available for discriminating CTE from co-morbid pathologies may lead to an underrepresentation of CTE based on routine neuropathological analyses.

Could all cases of LATE-NC be associated with CTE? At first glance, this seems unlikely. Even though a small number of CTE cases have no known history of head injury^45^, neuropathologic CTE is closely associated with repetitive head injury, particularly in the context of contact sports^29–31^. At the same time, no such relationship exists for LATE-NC. Moreover, CTE is rare in neurodegenerative brain banks^46^, whereas LATE-NC is exceedingly common^18–20^. Our study highlights the diagnostic challenges inherent to neuropathologic CTE, including the fact that pathognomonic cortical lesions required for diagnosis can be sparse and therefore missed without extensive sampling (which still represents <1% of cortex by volume)^32^. Furthermore, CTE lesions can resemble or be masked by ARTAG^47^. In addition, extracortical CTE pathology is difficult to distinguish from other mixed 3R/4R tauopathies accurately^32^. Consistent with this, CTE was not suspected based on the initial review of any of these cases; instead, diagnostic lesions or findings suggestive of CTE were identified only upon additional sampling prompted by our structural analysis.

Conversely, whether the TDP-43 pathology observed in CTE represents LATE-NC is unclear. TDP-43 pathology is a frequent component of CTE, found in ∼40% of cases^33^. In cases of “pure” CTE, TDP-43 pathology is confined to the cortex, brainstem, and/or spinal cord, and therefore is relatively distinct from LATE-NC, where it is concentrated in the medial temporal lobes^34,35^. Nevertheless, TDP-43 pathology in older patients with co-morbid CTE and AD displays a limbic-predominant distribution resembling that of LATE-NC, and is often associated with hippocampal sclerosis^33^. This is precisely what we observed in case 1, in which we confirmed the presence of CTE in addition to AD+LATE-NC. Whether TDP-43 pathology in CTE and LATE-NC reflects distinct processes or a pathologic continuum remains unclear. Further investigation into the incidence of CTE neuropathology in LATE-NC is urgently needed to help answer this question.

An alternative possibility is that LATE-NC is distinct from CTE yet driven by CTE-fold tau. Both SSPE and ALS/PDC lack pathognomonic CTE lesions, but exhibit NFTs concentrated in superficial cortical layers reminiscent of CTE^48–50^. Moreover, both SSPE and ALS/PDC are associated with i) TDP-43 pathology, and ii) antecedent brain insult/injury, much like CTE. SSPE occurs decades after measles infection and is presumed to be secondary to diffuse brain inflammation^48^. While the antecedent brain insult in ALS/PDC is unknown, it is likely to be environmental, given the limited geographic distribution of the disorder and the rapid decline in incidence upon modernization of Guam^51^.

These unifying features of CTE-fold tauopathies, together with our data illustrating CTE-fold tau in LATE-NC, hint at a fundamental relationship between CTE-fold tau and TDP-43 (**Fig. 4c**). In keeping with this, a growing body of evidence suggests that TDP-43 pathology influences the structural conformations of tau and downstream toxicity associated with these conformation^52–55^. Furthermore, our cryo-EM analysis revealed the presence of CTE-fold tau in all three cases of AD+LATE-NC that we analyzed, supporting the intriguing possibility that TDP-43 pathology in LATE-NC facilitates the formation of CTE-fold tau filaments. The connection we uncovered between CTE and LATE-NC also raises the question of brain injury as a potential risk factor for LATE-NC. Addressing these issues is critical if we are to successfully understand, identify, and treat these conditions. Still, it will require more accurate and generalizable means of identifying CTE fold-tau and CTE, particularly in the setting of co-morbid conditions featuring deposition of TDP-43 and other neurodegenerative disease-associated proteins.

## Author Contributions

E.S.P. identified all patient cases, obtained samples, and performed neuropathologic analysis together with K.S.C., and A.P.L. V.N. and M.G.F. established and optimized the protocols for filament isolation from patient tissues. J.K.R., E.S.P., and M.G.F. purified the tau filaments.

J.K.R. and S.M. prepared the cryo-EM grids. J.K.R. collected the cryo-EM data, performed image analysis, and solved the structures (PHF, SF, CTE) for all cases. J.K.R., E.S.P., and S.M. prepared the figures and wrote the first draft of the manuscript. A.P.L. and K.S.C. reviewed neuropathology for all three cases. All authors contributed to the final draft of the manuscript.

S.J.B. and S.M. supervised the project and provided all the resources for the study.

## Acknowledgments

This work was supported by: National Institutes of Health DP2GM150019-01, and Klatskin Sutker Discovery Fund (S.M.), National Institutes of Health R01NS097542, R37NS113943 and 1R56NS128110-01 (S.J.B), Kissick Family Foundation & Milken Institute Frontotemporal Dementia Grant (S.J.B. & S.M.), National Institutes of Health AWD012778 (E.P.), National Institutes of Health T32GM141840 to (M.G.F.), National Institutes of Health P30AG072931 to the University of Michigan Brain Bank and Alzheimer’s Disease Research Center, National Institutes of Health S10OD030275 and the Arnold and Mabel Beckman Foundation award to the University of Michigan Cryo-EM facility. The funders had no role in the study design, data collection, analysis, or the content and publication of this manuscript.

## Competing interests

S.J.B. serves on the advisory board for Neurocures, Inc., Symbiosis, Eikonizo Therapeutics, Ninesquare Therapeutics, the Live Like Lou Foundation, and the Robert Packard Center for ALS Research. S.J.B. has received research funding from Denali Therapeutics, Biogen, Inc., Lysoway Therapeutics, Amylyx Therapeutics, Acelot Therapeutics, Meira GTX, Inc., Prevail Therapeutics, Eikonizo Therapeutics, and Ninesquare Therapeutics.

## Data availability

Cryo-EM maps of tau SF, PHF, and CTE have been deposited in the Electron Microscopy Data Bank (EMDB) under accession codes EMD-73448, 73449, 73450 for case 1, EMD-73451, 73452, 73453 for case 2, and EMD-73454, 73455, 73456 for case 3, respectively. All other materials used and data reported in the study are available upon request.

