## Supplementary Material for "CTE-Type Tau Filaments in Alzheimer’s Disease with Co-morbid LATE-NC"

**This file contains:**

Methods

Extended Data Figures 1-7

### Methods

*Clinical history and neuropathology*

Individual 1 (Case 1) presented at age 82 with “leaning attacks” interpreted as transient ischemic attacks. At age 89, the individual began to show multi-domain cognitive impairment with difficulty navigating familiar places, word-finding difficulties, and trouble consistently taking medications. Cognition and motor function steadily declined until death at age 94. The clinical diagnosis, at death, was “mixed cognitive and motor disease”.

Individual 2 (Case 2) presented at age 69 with memory impairment and was given a clinical diagnosis of Alzheimer’s Disease. Cognition was relatively stable for the first few years, but then progressively declined until the individual’s death at age 80.

Individual 3 (Case 3) presented at age 74 with balance difficulties and frequent falls. Beginning around age 78, the individual developed cognitive symptoms, including forgetting names and faces. Motor symptoms became progressively more severe, and cognition was relatively stable until death at age 86. Clinical diagnosis at the time was “corticobasal degeneration”.

Extended clinical and pathological information for all individuals is presented in Table 1.

*Immunohistochemistry*

Detailed immunohistochemical analysis of different brain regions is provided in Tables 4-6 for cases 1-3, respectively. Immunohistochemistry was performed on a Ventana Benchmark Ultra immunostainer (Tucson, AZ). Slides were dewaxed, rehydrated, and subjected to antigen retrieval (details vary by antibody; see Table 2). Slides were incubated with the primary antibody and detected using the indicated system, as noted in Table 2. Slides were then counterstained with hematoxylin and coverslipped.

*Purification of filaments from patient tissue*

Sarkosyl-insoluble material was extracted from flash-frozen amygdala using protocols described previously[56]. Briefly, ~1.0 g of tissue was homogenized in 40 ml buffer A (10 mM Tris-HCl, pH 7.5, 0.8 M NaCl, 10% sucrose, and 1 mM EGTA) using a polytron homogenizer and solubilized in 2% final sarkosyl solution for 1 h at 37 °C. The homogenates were centrifuged at 27,000 *g* for 15 min, and the supernatant was collected for further centrifugation at 166,000 *g* for 30 min. The pellets were resuspended in 6 mL $g^{-1}$ buffer A containing 1% sarkosyl and homogenized using a hand-held probe sonicator. The homogenates were diluted 4-fold with buffer A containing 1% sarkosyl and incubated for an additional 30 min at 37 °C. The solubilized suspension was first centrifuged at 17,000 *g* for 5 min at 25 °C, followed by centrifugation at 166,000 *g* for 30 min at 25 °C. The resulting pellets were resuspended in 1 ml g-1 buffer A containing 1% sarkosyl and incubated for 1 h at 37 °C. Final centrifugation was performed at 100,000 *g* for 30 min at 25 °C. The sarkosyl-insoluble pellet was resuspended in 30 µl of buffer B (20 mM Tris-HCl, pH 7.4, and 150 mM NaCl).

*Immunoblotting analysis*

The tissue lysate (T), supernatant post-sarkosyl-solubilization (S), and the sarkosyl-insoluble pellet (I) were resuspended in a final working solution of 1× Laemmli buffer (Bio-Rad, #161-0747), heated at 95 °C for 5 min, resolved on 4-15% Criterion TGX$^{TM}$ (Bio-Rad) gels at 200 V for 40 min, and transferred onto PVDF membrane using Trans-Blot Turbo Transfer System (Bio-Rad). The membranes were blocked in PBS containing 1% Bovine Serum Albumin and 0.2% Tween-20 for 1 h at 21 °C. The membranes were subsequently incubated with primary antibodies: phospho-TDP-43 (Ser 409/410, 1:1000; Proteintech, #22309-1-AP), phospho-tau (Ser202/Thr205, 1:1000; Invitrogen AT8), Aß (1:1000, Milipore Sigma, #A3981), and TMEM106B[57] (1:800) overnight at 4 °C. The blots were washed three times with PBS containing 0.2% Tween-20 and subsequently incubated with HRP-conjugated secondary antibody, Goat anti-Rabbit (BD Biosciences, #554021) or Goat anti-Mouse (Millipore Sigma, 12-349) for 1 h at 21 °C. Following this, the membranes were again washed three times with PBS containing 0.2% Tween-20 and developed using Clarity Western ECL substrate (Bio-Rad, #1705061) and images using Bio-Rad ChemiDoc Imaging System.

*Immunoelectron microscopy (Immuno-EM)*

300 mesh carbon-coated copper grids (Electron Microscopy Sciences, #CF300-CU-50) were glow-discharged for 60 s at 5 mA (PELCO easiGlow). Sarksosyl-insoluble filaments were further diluted 1:10 in buffer B (20 mM Tris-HCl, pH 7.4, and 150 mM NaCl), and 3 µl of the sample was applied to the grids, incubated for 3 min, and subsequently the excess liquid was blotted from the grids using a Whatman 1 filter paper (Cytiva, #1001090). Next, blocking buffer [PBS, pH 7.4, 0.1% w/v cold water fish skin gelatin (Aurion, CFWS Gelatin)] was applied, and the grids incubated for 10 min. Subsequently, excess liquid was blotted again, and primary antibody (see immunoblotting analysis; 1:20 in blocking buffer) was applied to the grids and incubated for 1 h. Following blotting, grids were washed five times with a blocking buffer, and excess solution was removed in between each wash. 12 nm Colloidal Gold AffiniPure Goat Anti-Rabbit IgG (Jackson ImmunoResearch Laboratories, #111-205-144) diluted 1:4 in blocking buffer was applied to the grids and incubated for 1 h. Grids were washed five times with Milli-Q water, and excess liquid was blotted off between washes. Finally, grids were stained with 3 µL of 0.75% uranyl formate (Electron Microscopy Sciences, #16984-59-1) for 1 min, washed twice with Milli-Q water, and the excess solution blotted off. The grids were imaged using a Morgagni transmission electron microscope (Thermo Fisher Scientific) at an acceleration voltage of 100kV. The images were recorded with a Gatan Orius SC200 CCD camera with a resolution of 2.1 Å/pixel and Digital Micrograph software (Gatan Inc).

*Cryo-EM Sample preparation and data-collection*

Sarkosyl-insoluble pellets were diluted 1:4 in buffer B (20 mM Tris-HCl, pH 7.4, and 150 mM NaCl) and treated with 0.4 mg ml-1 pronase (Sigma) before plunge-freezing. 2.5 - 3 µl of samples were applied to glow-discharged (60 s at 15 mA, PELCO easiGlow) R1.2/1.3 UltrAuFoil Au 300 mesh grids (Quantifoil GmbH). Grids were blotted for 3.5 s at 18 °C and 100% humidity with blot force 5, then immediately frozen in liquid ethane using a Vitrobot (Mark IV, Thermo Fisher Scientific). Images were acquired using Titan Krios G4i operated at 300 kV equipped with a bioQuantum energy filter with a slit width of 20 eV and a Gatan K3 direct electron detector. Automated collection was performed at a nominal magnification of 105,000×, yielding a calibrated pixel size of 0.83 Å/pixel with SerialEM v 4.0[58]. A total of 50 frames were collected during a 2 s exposure, with an accumulated dose of 50.4 $e^- Å^{-2}$, and a defocus range of -2.5 µm to -1 µm.

*Helical Reconstruction*

All image processing was performed using RELION 5.0.0[59]. Movies were gain-corrected, aligned, dose-weighted, and summed using the motion-correction algorithm implemented in RELION-5.0.0[60]. Motion-corrected micrographs were used to estimate the contrast transfer function using CTFFIND4[61]. Helical reconstruction of tau filaments was performed as described previously[62–64]. Briefly, the filaments were manually picked and extracted as segments with a box size of 1024 pixels, then downsampled to 128 pixels. Two rounds of reference-free 2D classification were performed, and particles were selected based on the hierarchical clustering of the filaments using the FilamentTools program, resulting in a homogenous subset of the class averages for each filament type[43]. Particle numbers at this stage were used to calculate percentage ratios of the polymorphs depicted in Fig. 4a.

In each case, initial models were generated *de novo* using *relion_helix_inimodel2d.py* as previously described, to avoid any model bias. For initial model generation, a subset of particles for each filament type was re-extracted with a box size of 768 pixels, downsampled to 384, subjected to reference-free 2D classification, and used as inputs. The volumes obtained were rescaled and used as initial references for 3D auto-refinement, starting with 8× binned particles (6.64 Å/pixel) and progressing to an unbinned final refinement (0.83 Å/pixel). Optimization of helical parameters was performed once the map resolution was at 4.5Å or lower, and a clear separation among β-sheets was apparent. In all three cases, we achieved resolutions of 3.2-3.6 Å for all the structures. Bayesian polishing was performed to improve the resolution of each map further[60]. For only cases 1 and 2, 3D classification was performed to identify the best-aligned segments, followed by CTF refinement to improve map resolution further[65]. No 3D classification was performed for particles in case 3. Final 3D reconstructions from all cases (a total of 9 maps) were sharpened using the post-processing method implemented in RELION, and global resolution was estimated from Fourier shell correlation at a cutoff of 0.143 between the two independently refined half-maps. Post-processed maps were symmetrized with helical symmetry parameters in real space using the RELION Helix Toolbox program.

*Model building*

We build *de novo* models using ModelAngelo[66]. RMSD between the published structures and ModelAngelo models was less than 1. We therefore used published Protein Data Bank (PDB) models of PHF (PDB ID: 5O3L), SF (PDB ID: 5O3T), and CTE (PDB ID: 6NWP) to fit into the corresponding maps. For SF, PHF, and CTE, the published models were fit into corresponding Coloumbe potential maps as rigid bodies in ChimeraX[67]. After removing hydrogen atoms in COOT[68], real-space refinement was performed using PHENIX[28]. ChimeraX was used to visualize the structures and make figures.

**Extended Data Figures (1-7):**

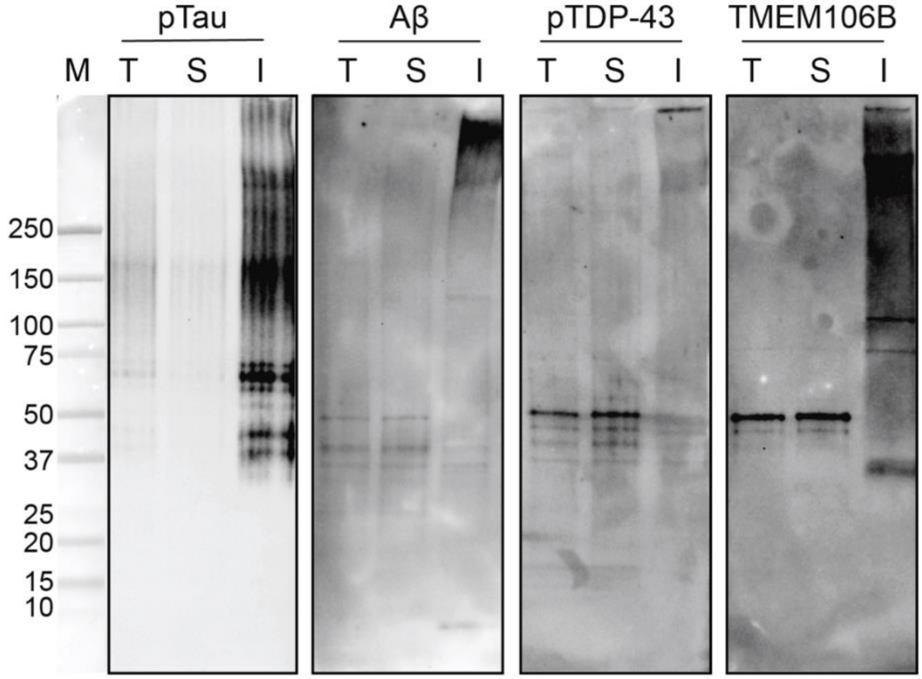


**Extended Data Figure 1: Immunoblot analysis of insoluble inclusions from the amygdala of a patient with LATE with AD (Case 1).** Sarkosyl-insoluble material (I) was isolated from the amygdala, and probed with antibodies AT8 (specific for pS202 and T205 tau; pTau), anti-amyloid beta antibody (Aβ), anti-pS409/S410 TDP-43 antibody (pTDP-43), or anti-TMEM106B core[57]. Total lysate (T) and the soluble fraction (S) are also shown in each case. The Precision Plus Protein Dual Color (Bio-Rad) marker is denoted as M.

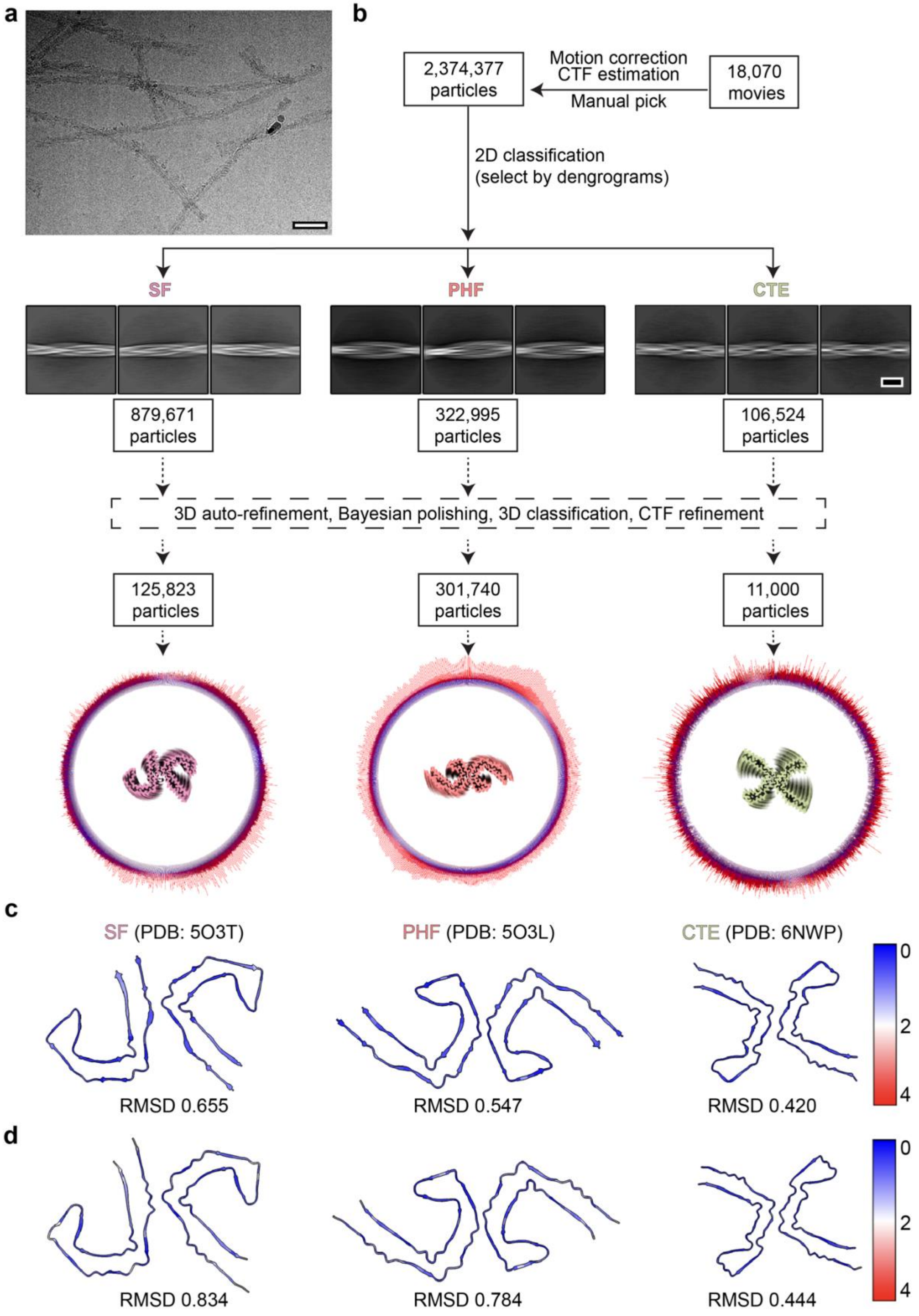

a
b
2,374,377 particles
Motion correction
CTF estimation
Manual pick
18,070 movies
2D classification
(select by dengrograms)
SF
PHF
CTE
879,671 particles
322,995 particles
106,524 particles
3D auto-refinement, Bayesian polishing, 3D classification, CTF refinement
125,823 particles
301,740 particles
11,000 particles
c
SF (PDB: 5O3T)
PHF (PDB: 5O3L)
CTE (PDB: 6NWP)
RMSD 0.655
RMSD 0.547
RMSD 0.420
0
2
4
d
RMSD 0.834
RMSD 0.784
RMSD 0.444
0
2
4

**Extended Data Figure 2: Helical reconstruction workflow for tau filaments. a.** Representative cryo-EM image of sarkosyl-insoluble material from amygdala. Scale bar, 50 nm. **b**. Data analysis workflow for the structure determination of tau filaments from case 1 as a representative example. ~18,000 movies were collected, motion corrected, and CTF was estimated on motion corrected micrographs using established procedures in Relion-5[60,61]. Fibrils were manually picked and sorted using dendrograms. The representative 2D classes are shown for SF, PHF, and CTE (scale bar, 10 nm). The filaments were separated, and all downstream steps were performed independently on each particle stack. **c**. Comparison between the model presented in the current study and the published model for SF (PDB: 5O3T), PHF (PDB: 5O3L), and CTE (PDB: 6NWP). The models are colored based on the RMSD values from 0 (blue), 2 (white), and 4 (red). **d**. Comparison between the SF, PHF, and CTE models determined by ModelAngelo[66] and the published model for SF, PHF, and CTE. The models are colored based on the RMSD values from 0 (blue), 2 (white), and 4 (red).

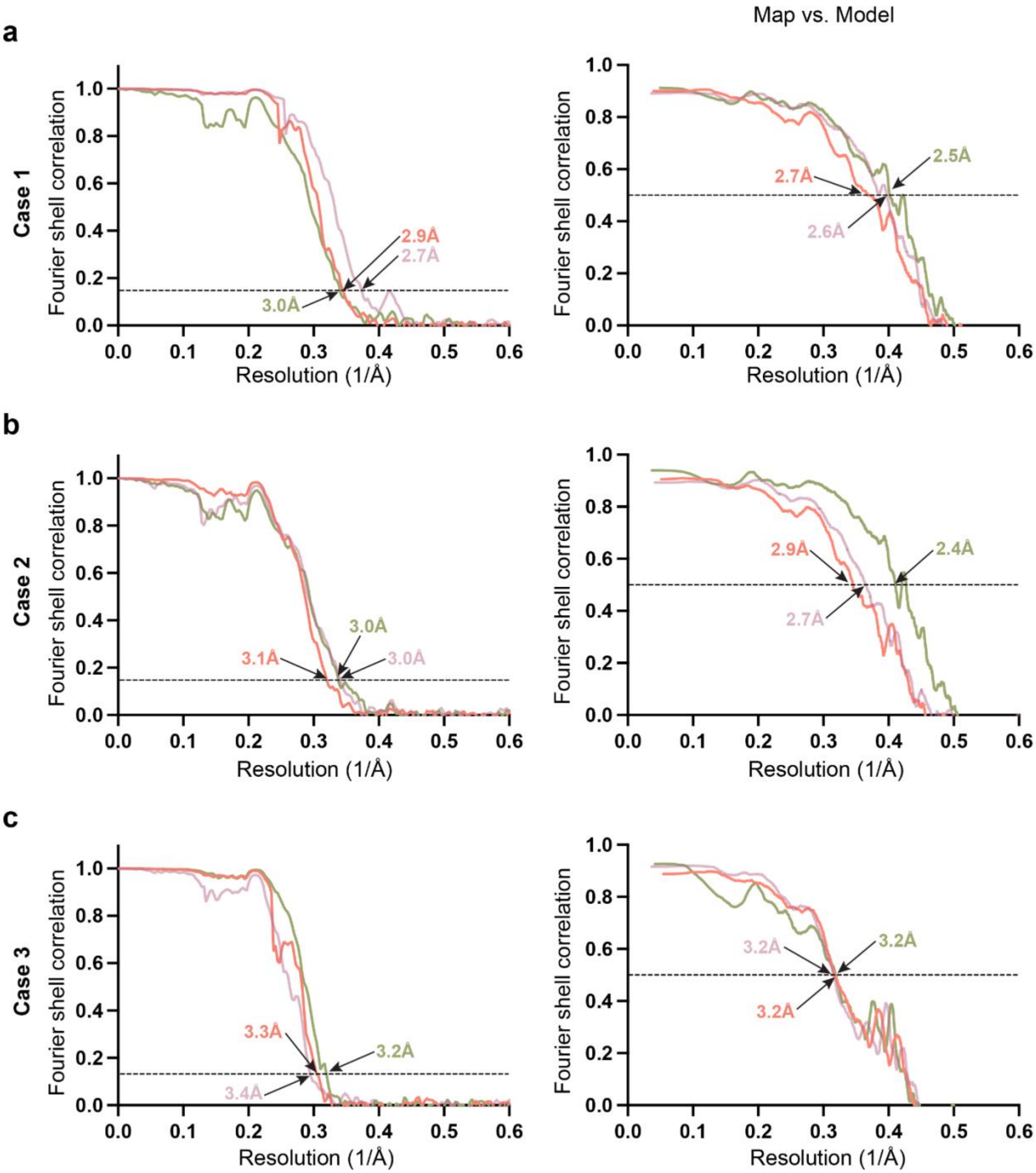


**Extended Data Figure 3: Fourier shell correlation (FSC) curves. a-c.** FSC curves of cryo-EM maps (left) and model to map validation (right) for all three LATE with AD cases (cases 1-3, respectively) analyzed in this manuscript. In each case, curves for SF, PHF, and CTE are colored in magenta, warm pink, and green, respectively. Resolution in angstroms is shown for the cryo-EM maps ($FSC_{0.143}$) and model to map ($FSC_{0.5}$).

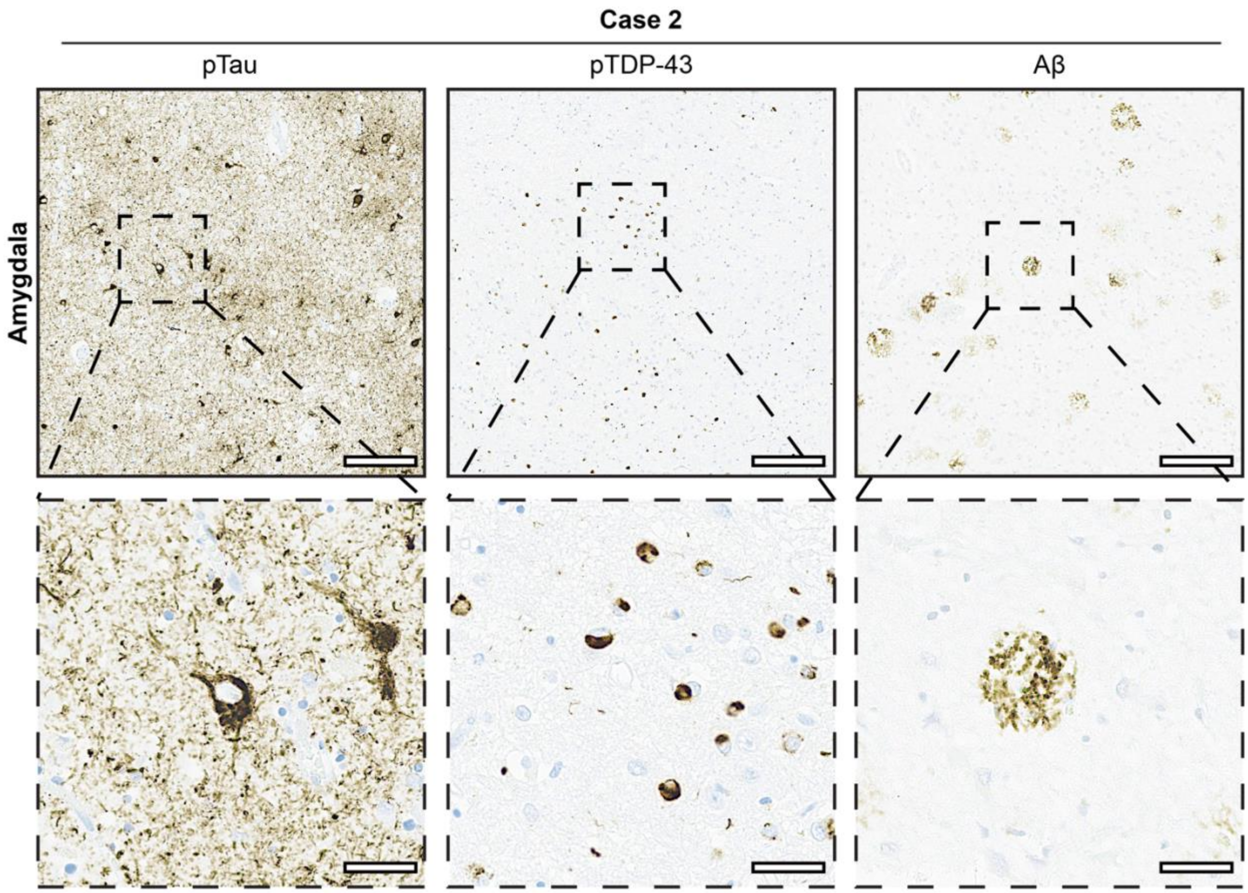


**Extended Data Figure 4: Immunohistochemical analysis of pathological inclusions in the amygdala of a patient with LATE with co-morbid AD. Case 2.** Representative images of amygdala immunostained with AT8 (specific for pS202 and T205 tau; pTau, top left), anti-pS409/S410 TDP-43 antibody (pTDP-43, middle), and anti-amyloid beta antibody (Aβ, top right). Insets show indicated high magnification views. Scale bars: 300 μm (top) and 60 μm (bottom).

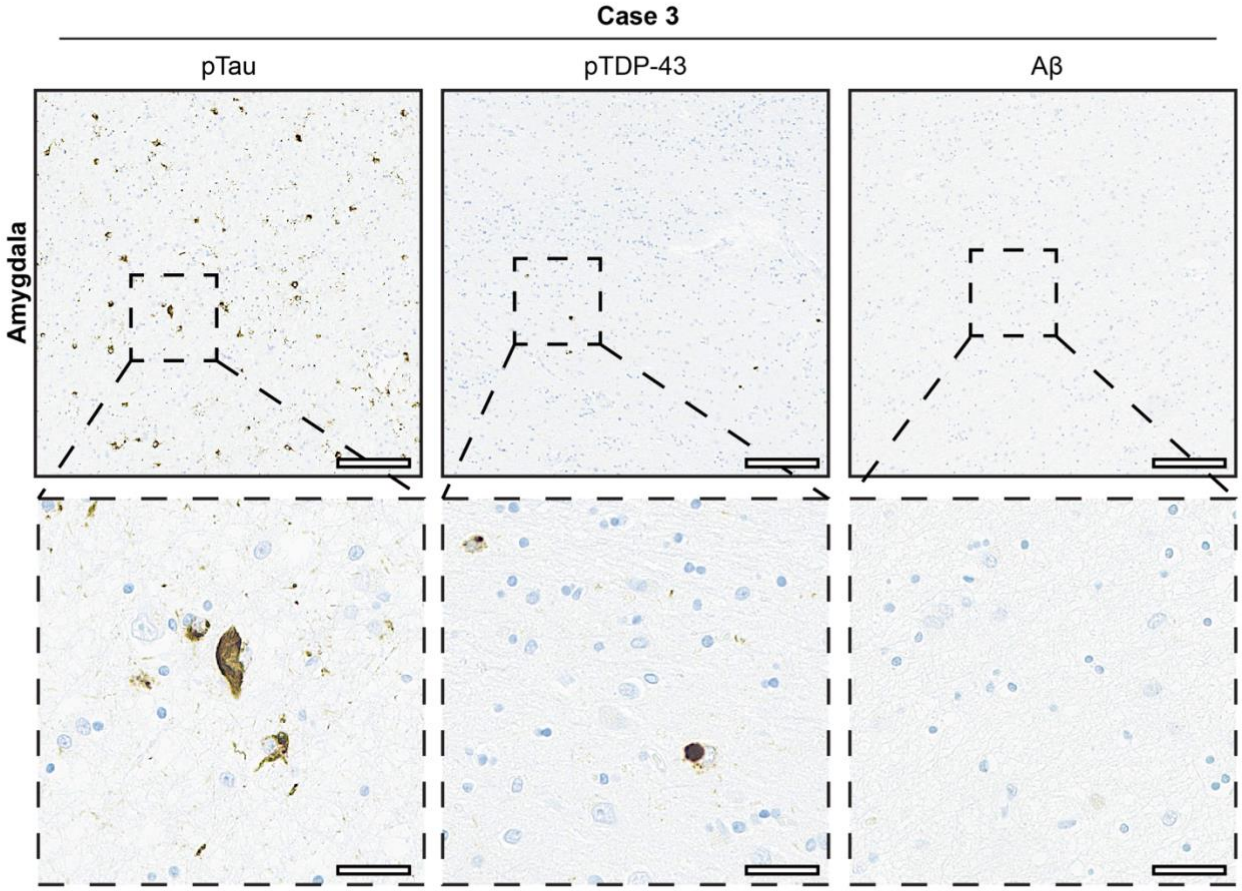


**Extended Data Figure 5: Immunohistochemical analysis of pathological inclusions in the amygdala of a patient with LATE with co-morbid AD. Case 3.** Representative images of amygdala immunostained with AT8 (specific for pS202 and T205 tau; pTau, top left), anti-pS409/S410 TDP-43 antibody (pTDP-43, middle), and anti-amyloid beta antibody (Aβ, top right). Insets show indicated high magnification views. Scale bars: 300 μm (top) and 60 μm (bottom).

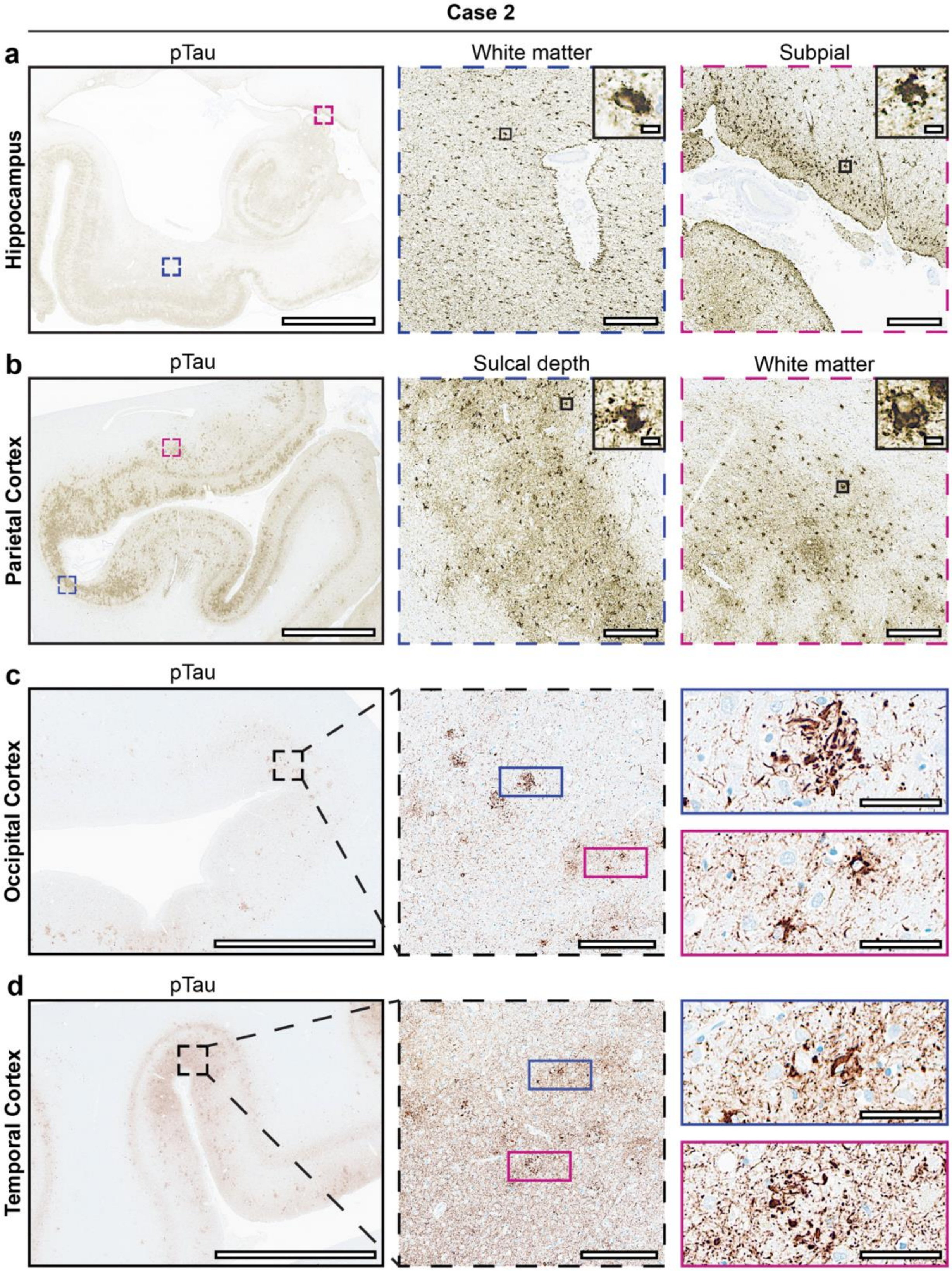


**Extended Data Figure 6: Widespread aging-related tau astrogliopathy (ARTAG) and suggestive features of CTE in Case 2.** Representative AT8 (specific for pS202 and T205 tau;

pTau) immunostaining in hippocampus (**a**), parietal cortex (**b**), occipital cortex (**c**), and temporal cortex (**d**). Insets show indicated high magnification views. Scale bars: 5mm (left a-d), 250 μm (center and right a-b, middle c-d), 10 μm (insets a-b), and 50 μm (right c-d).

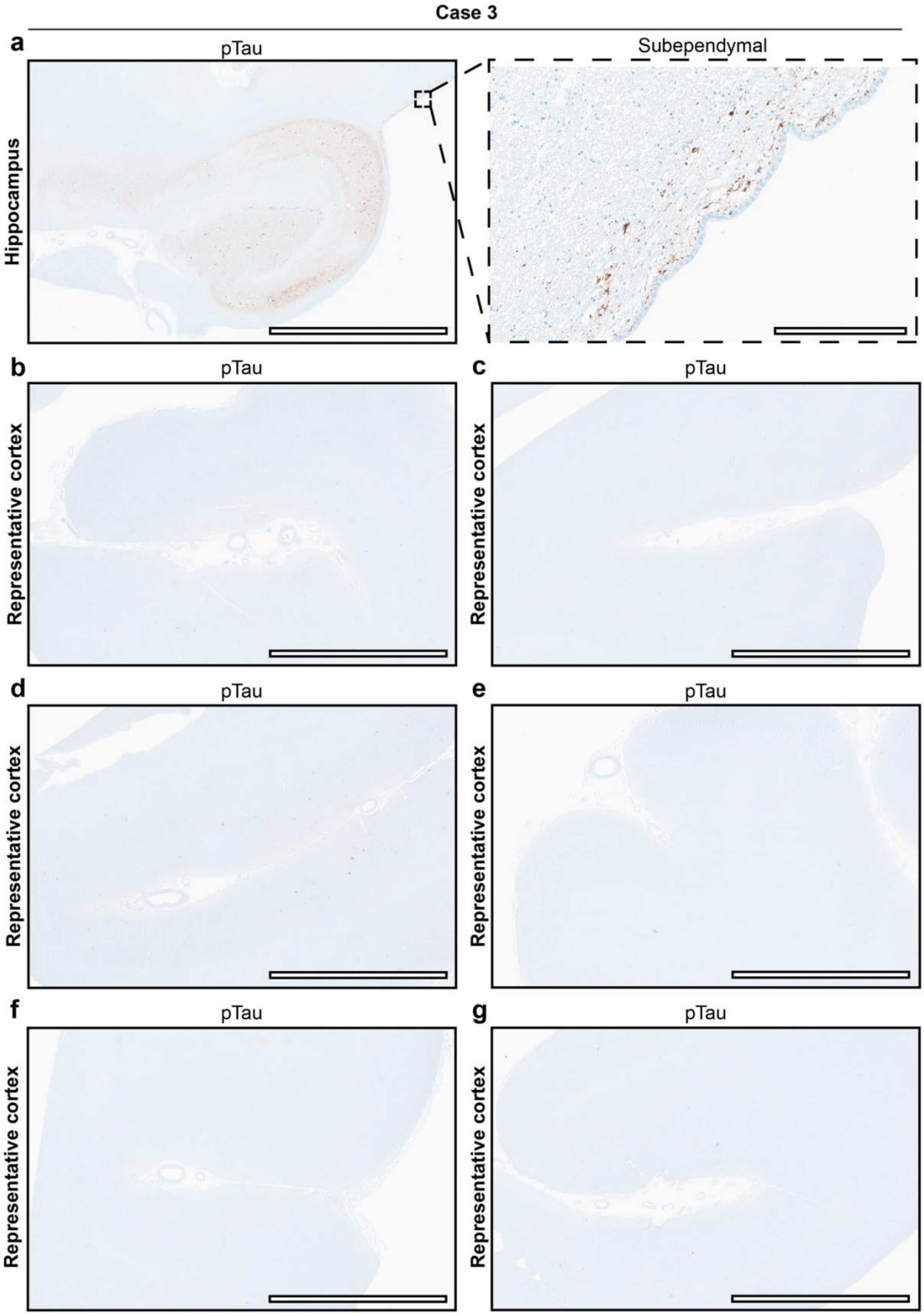
Case 3
a
pTau
Subependymal
Hippocampus
b
pTau
Representative cortex
c
pTau
Representative cortex
d
pTau
Representative cortex
e
pTau
Representative cortex
f
pTau
Representative cortex
g
pTau
Representative cortex

**Extended Data Figure 7: Focal ARTAG and no evidence of CTE in Case 3.** Representative AT8 (specific for pS202 and T205 tau; pTau) immunostaining in hippocampus (**a**) and representative cortical sections (**b-g**). The inset shows indicated high magnification views. Scale bars: 5mm (left a and b-g) and 250 μm (right a).
